# Selection for antibiotic resistance is reduced when embedded in a natural microbial community

**DOI:** 10.1101/529651

**Authors:** Uli Klümper, Mario Recker, Lihong Zhang, Xiaole Yin, Tong Zhang, Angus Buckling, William Gaze

## Abstract

Antibiotic resistance has emerged as one of the most pressing, global threats to public health. In single-species experiments selection for antibiotic resistance occurs at very low antibiotic concentrations. However, it is unclear how far these findings can be extrapolated to natural environments, where species are embedded within complex communities. We competed isogenic strains of *Escherichia coli*, differing exclusively in a single chromosomal resistance determinant, in the presence and absence of a pig fecal microbial community across a gradient of antibiotic concentration for two relevant antibiotics: gentamicin and kanamycin. We show that the minimal selective concentration was increased by more than one order of magnitude for both antibiotics when embedded in the community. We identified two general mechanisms were responsible for the increase in minimal selective concentration: an increase in the cost of resistance and a protective effect of the community for the susceptible phenotype. These findings have implications for our understanding of the evolution and selection of antibiotic resistance, and can inform future risk assessment efforts on antibiotic concentrations.

## Introduction

The emergence and spread of antimicrobial resistance (AMR) genes in bacterial pathogens has been identified as one of the major threats to human health by the World Health Organisation (WHO, 2014). Whilst AMR genes have been detected in ancient permafrost samples (D’Costa et al., 2011), anthropogenic use of antibiotics has caused a rapid increase in their prevalence (Knapp et al., 2010). A large body of theory and *in vitro* work has identified the role of ecological context, such as treatment regime and environmental heterogeneity, in AMR gene dynamics (Drlica, 2003; Drlica and Zhao, 2007; Gullberg et al., 2014, 2011). However, the majority of this work has not explicitly considered a crucial feature of microbial ecology: microbes are typically embedded within complex communities of interacting species. This is always the case within human and livestock microbiomes, in which antibiotic-imposed selection is likely to be particularly strong (Carlet, 2012). Here, we combine experiments and theory to determine how selection for AMR is influenced by the presence of other species derived from a natural gut microbial community. The focus of this study is selection for pre-existing resistance genes within a focal species, rather than selection on *de novo* variation arising through spontaneous mutations or acquired through horizontal gene transfer from another species.

Recent experimental studies suggest that selection for AMR genes in complex communities is occurring at antibiotic concentrations (the minimum selective concentration; MSC) that are much lower than those that prevent the growth of susceptible bacteria (minimum inhibitory concentration; MIC) (Lundström et al., 2016; Murray et al., 2018); as has been previously shown within single species *in vitro* (Gullberg et al., 2014, 2011; Liu et al., 2011). However, it is unclear how the presence of other microbial species affects the MSC. While the precise effect of other species is likely context-dependent, we hypothesise that the presence of the community will typically increase the MSC. Studies of single species suggest that resistant cells can afford protection to susceptible ones, through both, intracellular and extracellular degradation of antibiotics (Medaney et al., 2016; Sorg et al., 2016; Yurtsev et al., 2013), thus increasing relative fitness of susceptible strains and hence the MSC. However, excreted metabolites can both potentiate or decrease antibiotic efficacy, thus decreasing or increasing MSCs (Cao et al., 2012; Churski et al., 2012). Further, any costs associated with AMR may be enhanced by increased competition for resources, as, for example, has been observed with respect to resistance in flies to parasitoids (Kraaijeveld et al., 2001) and bacteria to viruses (Gómez and Buckling, 2011).

To explore the potential effects of community context on AMR selection, we competed isogenic *Escherichia coli* MG1655 strains, differing exclusively in a single chromosomal resistance determinant, in the presence and absence of a microbial community across a gradient of two different aminoglycoside antibiotics, kanamycin and gentamicin. We embedded the *E. coli*, commonly found in the anaerobic digestive tract of warm-blooded mammals (Tenaillon et al., 2010), within a pig faecal community in experimental anaerobic digesters in an attempt to partially mimic a gut environment. We additionally employed metagenomic analysis, community typing (16S) and mathematical modelling to provide insights into mechanisms underpinning community effects on AMR selection.

## Results

### Community context affects selection for gentamicin resistance

Isogenic strains of the focal species *E. coli*, with and without gentamicin resistance, were competed in the presence and absence of a pig faecal community across a 5 orders of magnitude gradient of gentamicin concentrations. Independent of antibiotic concentration the focal species increased in abundance during the 3 day evolution experiment from ~10% at inoculation to above 90% relative abundance based on 16S sequencing (Fig S1A&B). Both resistant and sensitive strains showed positive growth across the whole range of concentrations and both treatments with cell counts increasing by 2.25 to 3.96 orders of magnitude per day (Fig 1A).

**Fig 1.**
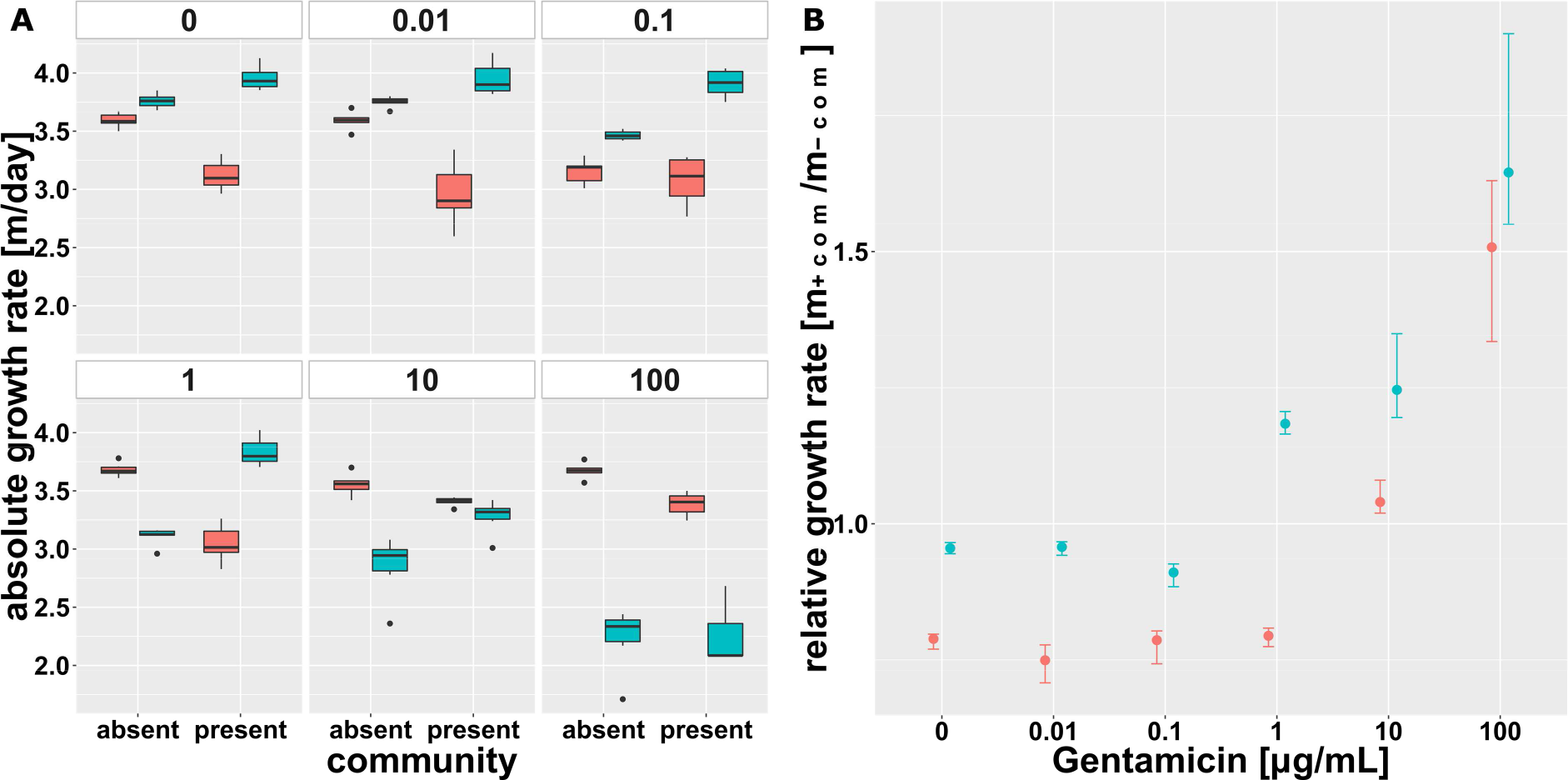
Malthusian growth parameter per day of the focal species’ isogenic strains for gentamicin. Values are displayed across the antibiotic gradient and in absence and presence of the gut microbial community. (A) Average (±SD, n=6) logarithmic absolute growth per day for the resistant (red) and the susceptible (blue) strain. Note: A different inoculum size of the focal species in absence (~10^6^ bacteria) and presence (~10^5^ bacteria = 10% of total inoculum) of the community was used. (B) Ratio of absolute Malthusian growth parameters (with 95% confidence intervals based on 1000-fold bootstrap analysis) in presence and absence of the microbial community across the gradient of antibiotic concentrations.

There was a small competitive fitness cost (t-test against 1, p=0.0005) of gentamicin resistance in the absence of the community (*ρ*_r_ = 0.955 ± 0.014, mean ± SD), and this cost appeared to be greatly increased when the community was present (Fig 1B, 2) (*ρ*_r_ = 0.788 ± 0.016) (ANOVA corrected for multiple testing, p<0.01, F=360.36). As antibiotic concentration increased, this cost was offset by the benefit of resistance. However, the reduction in fitness of the resistant strain in the presence of the community remained fairly constant (significant differences, p<0.05 after controlling for multiple testing at concentrations between 0 and 10 μg/mL) up to 100 μg/mL gentamicin, at which point the community had no effect on relative fitness (ANOVA corrected for multiple testing, p=0.259, F=1.42).

**Fig 2.**
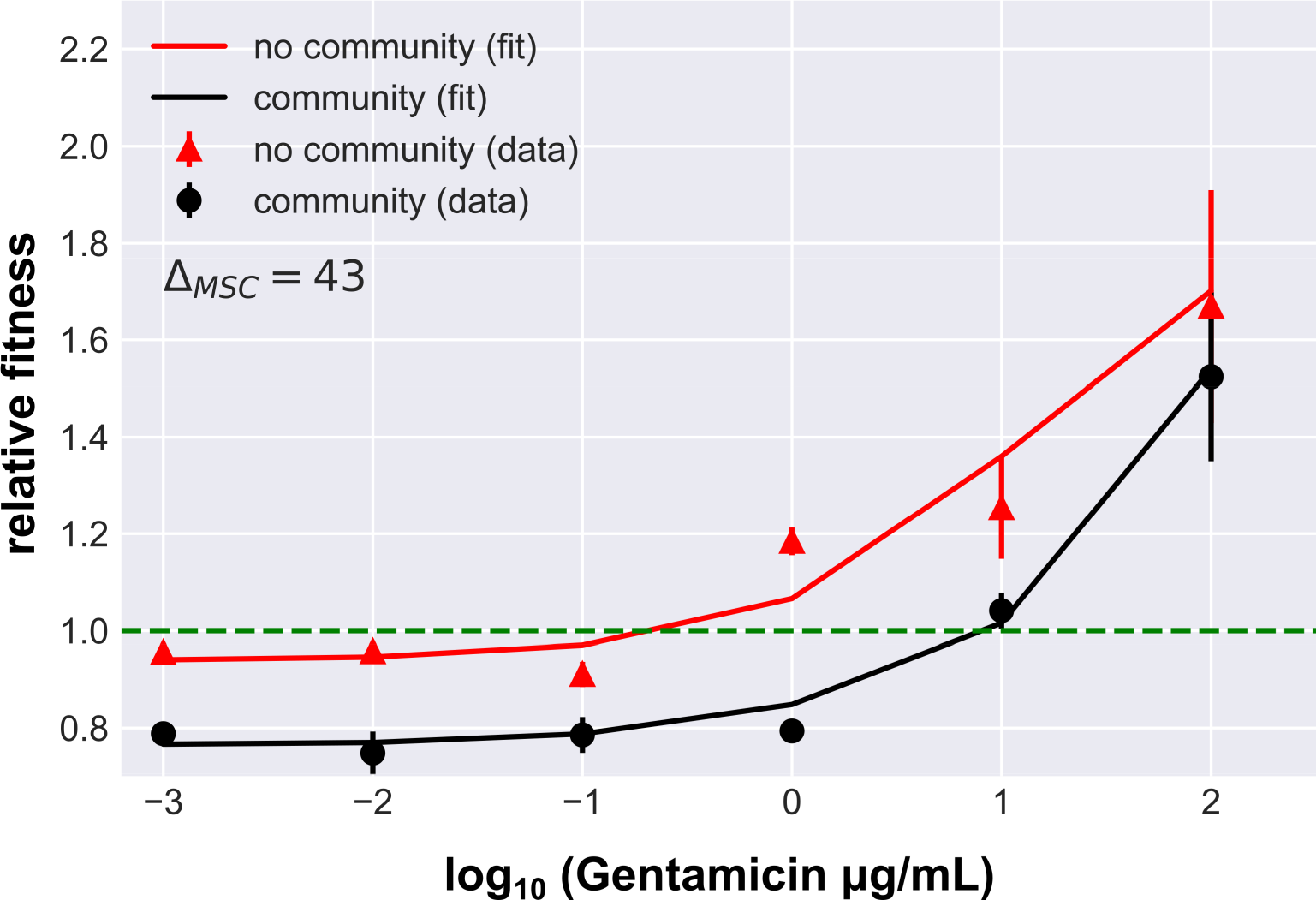
Relative fitness of the gentamicin resistant strain. Values (mean ± SD, n=6) in presence (black) and absence (red) of the community. Solid lines represent the best fit fitness curve through the mathematical model based on parameter estimates presented in Table 1. The dashed line indicates neutral selection at a relative fitness of *ρ*_r_ = 1, where the intercept with the fitness curve indicates the minimal selective concentration.

**Table 1:** Model parameter values for gentamicin fitness curves.

| parameter | Gentamicin |  |  |  |
| --- | --- | --- | --- | --- |
|  | all | susceptible | resistant | community |
| $\phi$ | | 1.4 | 1.3 | 1.3 |
| $e_{ij}$ | | 1 | 2.3 | 3 |
| $\alpha_{i,0}$ | | 1.3 | 2.9 | 1.6 |
| $\beta_{i,0}$ | | 0.7 | 0.8 | 0.6 |
| $k_d$ | $10^5$ | - | - | - |
| $f_{\max}$ | 0.9 | - | - | - |

#### Community composition is altered across the gentamicin gradient

It is possible that changes in community composition across the antibiotic gradient may have contributed to the observed changes in selection for resistance caused by the community, notably between 10 and 100 μg/mL. The composition of the microbial community changed significantly from the collected faecal sample, to inoculum and further during the duration of the experiment (AMOVA, p<0.001, Fig S2A). Above 1 μg/mL gentamicin the previously dominant Proteobacteria were outcompeted by Firmicutes (Fig S2B) leading to a significant (AMOVA, p<0.01) separation of communities below and above this threshold concentration in the NMDS plot (Fig S2B). However, there was no significant change in composition between 10 and 100 μg/mL, suggesting that compositional changes did not play a major role in community-imposed selection.

### Community context imposes a cost of resistance

To test the hypotheses derived from the numerical data we used numerical simulations of our experimental set up to determine the likely mechanisms underpinning the observed population dynamics in a common logarithmic growth model. We determined models based on the key empirical findings in the absence of the community (specifically, that there is a cost of resistance in the absence of antibiotics, and that antibiotics inhibit the growth of the sensitive strain in a dose dependent manner), and then determined the most parsimonious way in which the community could have altered the relative fitness of the resistant and susceptible strains (Table 1). We found a good fit to the data simply by assuming that the community imposed a greater competitive effect, constant across the antibiotic gradient, on the resistant rather than the sensitive strain (*e*_*rj*_ ≫ *e*_*cj*_ > *e*_*sj*_; where *e*_*ij*_ is the competition coefficient imposed on the focal population (resistant *r*, susceptible *s* and community *c*) by the community). Note that the lack of effect of the community at high antibiotic concentrations (100 μg/mL) was caused by there being very little growth of the susceptible strain, and hence relative fitness was determined primarily by the growth of the resistant strain.

The numerical simulation allowed us to estimate the change in MSC from absence to presence of the community by deterministically evaluating the concentration at the intercept with neutral selection at a relative fitness of *ρ*_r_ = 1. We estimated a 43-fold increase in MSC in the presence of the community (Fig 2).

### Community context affects selection for kanamycin resistance

As with gentamicin, the focal species increased in abundance during the 3 day competition experiment from ~10% at inoculation to above 90% relative abundance (Fig S1C&D). Again, both strains increased in abundance across both treatments and all concentrations of the 5 orders of magnitude antibiotic gradient with cell numbers increasing by 1.45 to 3.09 orders of magnitude per day (Fig 3A). In the absence of this community, kanamycin resistance also imposed a slight metabolic fitness cost on the resistant strain (*ρ*_r_ = 0.915 ± 0.036) (Fig 3B).

**Fig 3.**
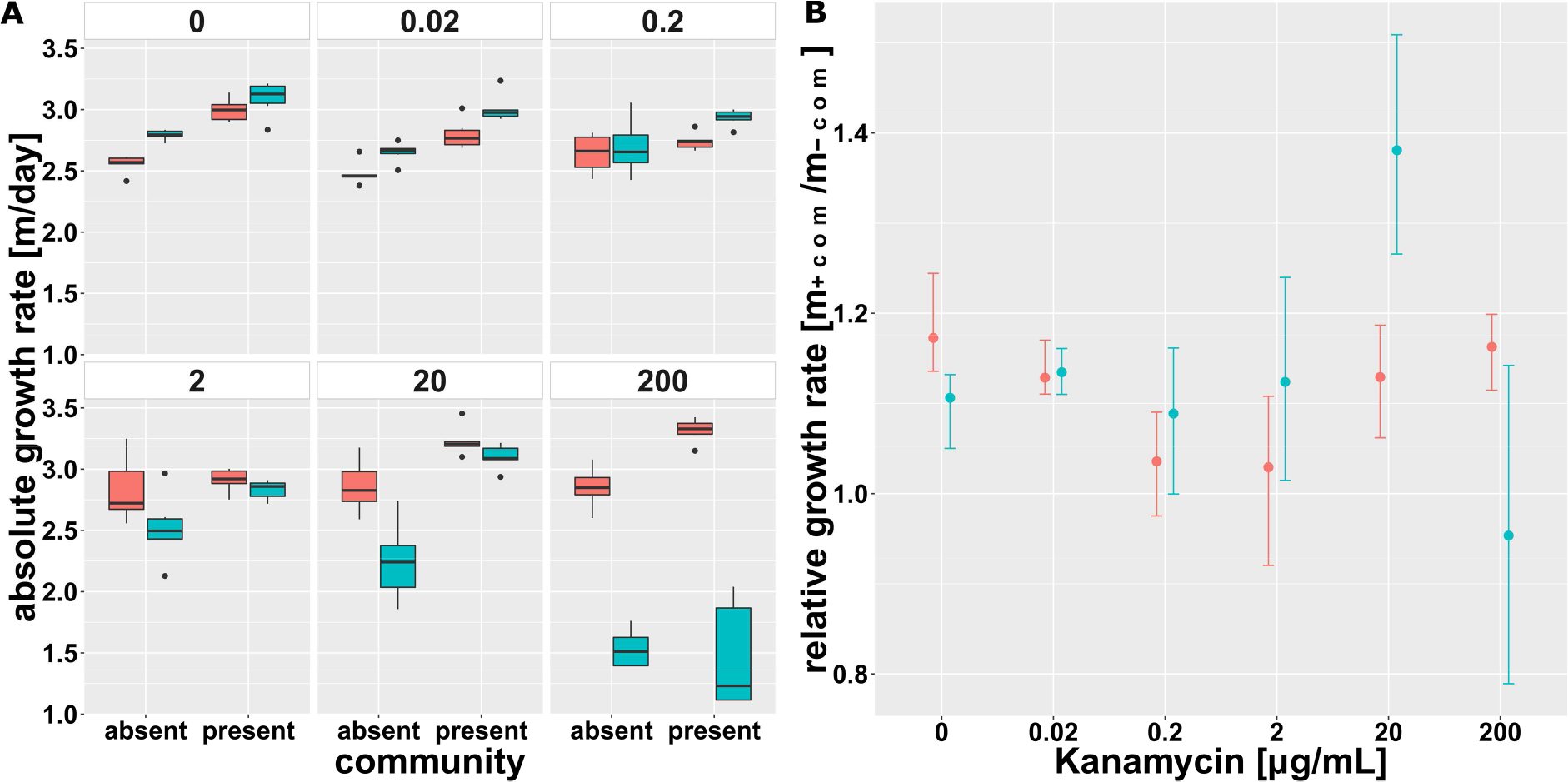
Malthusian growth parameter per day of the focal species’ isogenic strains for kanamycin. Values are displayed across the antibiotic gradient and in absence and presence of the gut microbial community. (A) Average (±SD, n=6) logarithmic absolute growth per day for the resistant (red) and the susceptible (blue) strain. Note: A different inoculum size of the focal species in absence (~10^6^ bacteria) and presence (~10^5^ bacteria = 10% of total inoculum) of the community was used. (B) Ratio of absolute Malthusian growth parameters (with 95% confidence intervals based on 1000-fold bootstrap analysis) in presence and absence of the microbial community across the gradient of antibiotic concentrations.

However, unlike gentamicin, the community did not increase the general cost of resistance. Indeed, the community had no significant effect on the relative fitness of the resistant strain except at a concentration of 20μg/mL (ANOVA corrected for multiple testing, p=0.002, F=15.58) (Fig 4). There was a clear fitness advantage for the resistant strain in the absence of the community at this concentration (*ρ*_r_ = 1.288 ± 0.149; t-test against 1, p=0.0052), while in the presence of the community this difference in relative fitness while still significant (t-test against 1, p=0.0088) was considerably lower (*ρ*_r_ = 1.034 ± 0.020). At 200 μg/mL kanamycin, close to the susceptible strains MIC, the resistant strain had an equally high relative fitness regardless of the presence of the community (ANOVA corrected for multiple testing, p=0.079, F=3.84).

**Fig 4.**
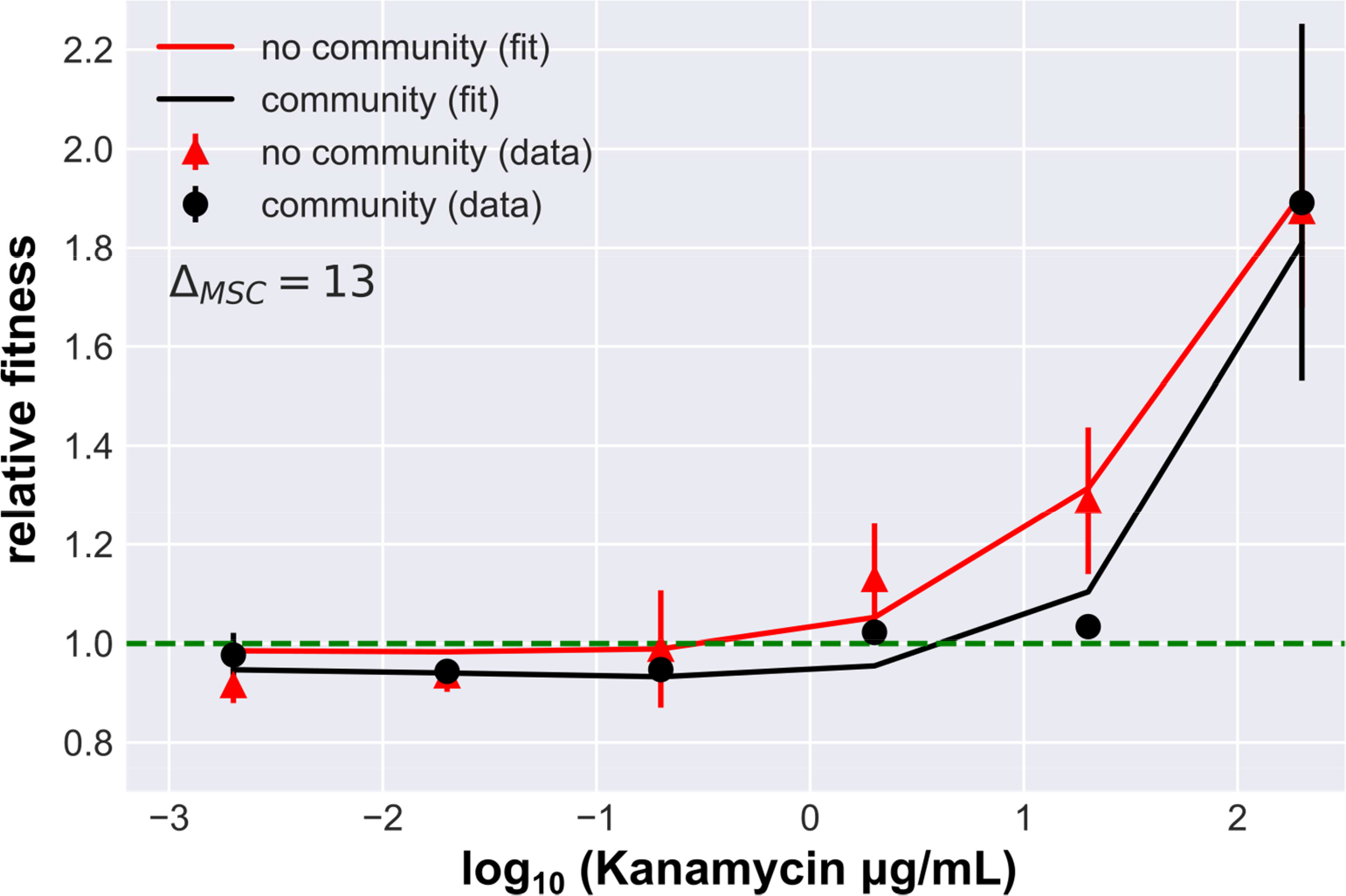
Relative fitness of the kanamycin resistant strain. Values (mean ± SD, n=6) in presence (black) and absence (red) of the community. Solid lines represent the best fit fitness curve through the mathematical model based on parameter estimates presented in Table 2. The dashed line indicates neutral selection at a relative fitness of *ρ*_r_ = 1, where the intercept with the fitness curve indicates the minimal selective concentration.

**Table 2.** Model parameter values for kanamycin selection curve.

| Kanamycin |  |  |  |  |
| --- | --- | --- | --- | --- |
| parameter | all | susceptible | resistant | community |
| $\varphi$ | - | 1.9 | 1.8 | 1.3 |
| $e_{ij}$ | - | 1.7 | 1.3 | 1.6 |
| $\alpha_{i,0}$ | - | 1.0 | 1.4 | 1.6 |
| $\beta_{i,0}$ | - | 0.6 | 0.4 | 0.5 |
| $k_d$ | $10^5$ | - | - | - |
| $f_{\max}$ | 0.9 | - | - | - |

#### Community and antibiotic resistance composition remain stable across the kanamycin gradient

As with the gentamicin experiment, a significantly shift in community composition from collected faecal sample, to inoculum and further during the duration of the kanamycin experiment (AMOVA, p<0.001, Fig S3A) was observed. However, across the whole gradient of antibiotics, Firmicutes (Fig S3B) remained the dominant phylum with no significant changes in community composition as a result. As such, compositional changes again cannot explain the impact of the community on focal strain fitness under selection at 20 μg/mL only. We additionally carried out metagenomic analysis for the 0, 2 and 20 μg/mL kanamycin treatments to determine whether relative abundance of resistance genes had changed within the community, despite the fact that there were no changes in community composition. Resistance to aminoglycoside (ANOVA, p=0.04) and other classes of antibiotics (fosmidomycin, kasugamycin, macrolides, polymyxin and tetracycline (ANOVA, all p<0.01)) significantly increased in the community of all reactors compared to the original faecal community independent of antibiotic concentrations (Fig 5A). However, there was no significant difference between kanamycin concentration and the abundance (ANOVA, p=0.15) of aminoglycoside resistance in general (Fig 5A) or any specific aminoglycoside resistance subtypes (Fig 5B) suggesting that relative fitness of the focal species was not influenced by the community resistome. Unsurprisingly, since no antibiotic concentration dependent selection for aminoglycoside resistance was observed within the community, no significant co-selection for resistance to any other classes of antibiotics was observed either.

**Fig 5.**
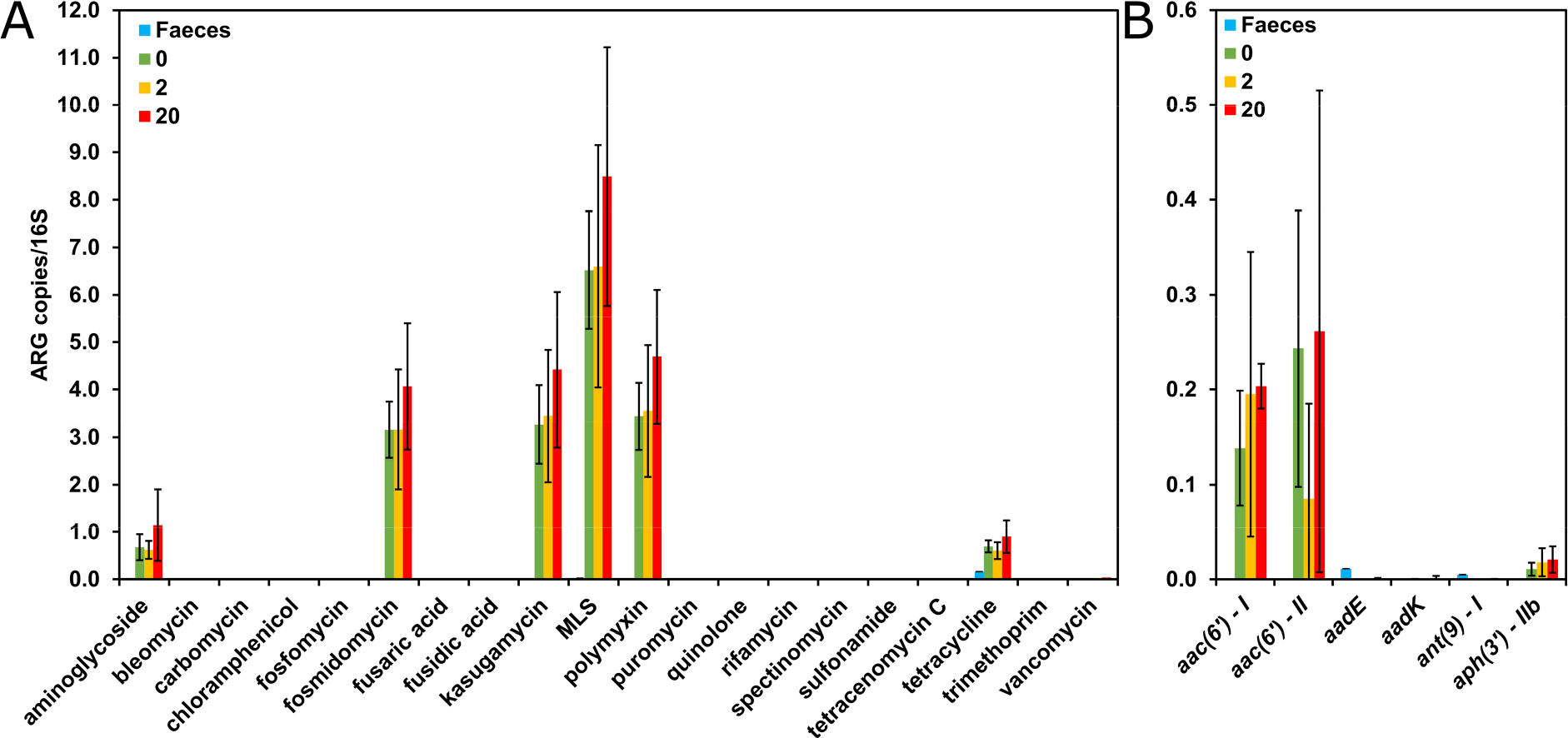
Detected resistance genes. Type (A) and aminoglycoside subtype (B) relative abundance (resistance gene number normalized with 16S rRNA copy number), in original faecal community and in final reactor community at 3 kanamycin concentrations (mean ± SD; n_feces_=2, n_Kn0_=6, n_Kn2_=6 n_Kn20_=5). Only genes detected with the ARGs-OAP pipeline are shown. MLS = Macrolides, Lincosamides, Streptogramines

#### Presence of the community can enhance growth of susceptible E. coli population at intermediate antibiotic concentrations

Numerical simulations showed that, unlike for gentamicin resistance, a community-imposed increase in the cost of kanamycin resistance was unable to explain why the benefit to the drug resistant focal *E. coli* strain was reduced in the presence of the community at intermediate drug concentrations (*e*_*rj*_ = *e*_*s*_). This suggested different interactions between *E. coli* and the rest of the community, and we speculated that the community might have provided a protective effect against kanamycin for the susceptible *E. coli*. Growth data demonstrated this to be the case: the presence of the community decreased or had no significant impact on the growth rate of either the susceptible or resistant *E. coli*, except at 20 μg/mL where the growth rate of the susceptible, but not the resistant strain, was significantly increased by the presence of the community (Fig 3B). We investigated if a protective effect of the community was sufficient to explain the observed data by fitting numerical simulations where the dose-response parameters *α*_*s*,*r*_ and *β*_*s*,*r*_ were explicitly dependent on the (time-dependent) density of the community (as listed in Table 2). The resulting model provided a good fit to the experimental data, suggesting that community protection was driving the observed population dynamics with a 12-fold increase in MSC.

## Discussion

In this study we investigated how being embedded within a semi-natural community (a pig gut derived community in an anaerobic digester) affects selection for AMR within a focal species (*E. coli*). For two antibiotics commonly fed to agricultural animals (gentamicin and kanamycin), we find the presence of the community selects against resistance, resulting in 1-2 orders of magnitude higher minimal selective concentrations for antibiotic resistance. This suggests that recent *in vitro* single strain based estimates of MSCs (Gullberg et al., 2014, 2011; Liu et al., 2011) are likely much lower than would be observed *in vivo* and might explain why in certain ecosystems no selection for antibiotic resistance was observed in focal strains (Flach et al., 2018).

The primary mechanisms responsible for this community-imposed reduction in selection for resistance differed for the two tested drugs, yet are likely fairly general based on their ecological origin. For gentamicin, the community increased the fitness costs reflected by reduced growth rates that are associated with resistance in the absence of antibiotics. These elevated costs were retained at similar levels across the antibiotic gradient, up until doses were so high that only the resistant strain grew (similar behaviour above a certain threshold concentration has previously been described for single strain systems (Andersson and Hughes, 2011, 2010) and our results show that his holds true in a community context). Resource limitation – directly manipulated or though competition - has been found to increase costs against a range of stressors in a range of organism, from resistance of plasmodium to antimalarial drugs (Wale et al., 2017) to phage resistance in bacteria (Gómez and Buckling, 2011). This is presumably because resource limitation has a more pronounced effect on resistant genotypes (Song et al., 2014).

For kanamycin, community-imposed selection against resistance was only apparent at intermediate antibiotic concentrations. The absolute growth rate of the susceptible strain was significantly increased at intermediate concentrations in presence of the community. Our model fitting suggests this is because of a protective effect of the community. The protective effect might have only been observed at intermediate concentrations since low concentrations were insufficient to detectably lower the relative fitness of the susceptible strain, while at high concentrations the protective effect was too small to be detectable. Such protective effects have been reported extensively within-species (Medaney et al., 2016; Yurtsev et al., 2013), as well as more recently within more complex communities (Sorg et al., 2016), either because of extra- or intracellular modification of antibiotics. Other common mechanisms known to increase a strains resistance to antibiotics in communities involve flocculation (Kümmerer, 2009) or biofilm formation (Drenkard and Ausubel, 2002; Mah et al., 2003), but might here only play a minor role due to the shaking conditions.

The mechanisms discussed above all underlie the selection for standing variation in pre-existing resistance genes, rather than selection on *de novo* variation arising through spontaneous mutations or horizontal gene transfer from other species. For de novo chromosomal mutations, the community is likely to further limit the spread of resistance, because the reduced population sizes of the focal strains in the presence of the community increases the chance that more costly mutations will be fixed (Perron et al., 2007). In contrast, being embed in a community might enhance the spread of resistance. First, there will be a greater source of resistance genes available to the focal species. Second, selection against resistance acquired through horizontal gene transfer at low antibiotic concentrations might follow different dynamics. While chromosomal resistance might be outcompeted and subsequently lost, resistance genes embedded on conjugative plasmids can persist or even increase in abundance, as a consequence of their sometimes extremely broad host ranges and high transfer frequencies (Arias-Andres et al., 2018; Klümper et al., 2017, 2015; Musovic et al., 2014; Shintani et al., 2014). In controlled single strain experiments plasmid born resistance proved more costly than chromosomal resistance (Gullberg et al., 2014). However, in more complex scenarios selection for mobile genetic element borne resistance usually depends not only on the single acquired resistance gene, but a combination of other linked traits encoded by the MGE as part of the communal gene pool (Norman et al., 2009). Thus, difficulties in making general predictions on the selection dynamics of horizontally acquired resistance in microbial communities arise that merit future research efforts.

In summary, we show that selection for antimicrobial resistance was influenced by being embed in a “natural” microbial community, such that the MSC was increased by more than one order of magnitude for two different antibiotics. Further to reducing relative fitness of resistance, being embedded in a community would also reduce absolute fitness, which has been argued to sometime be the major driver of spread of resistance (Day et al., 2015).

To determine MSCs that are relevant in environmental settings it is thus crucial to test for selection in a complex community context, rather than in single strain systems. Understanding under which concentrations selection for and thus long-term fixation of newly acquired resistance mechanisms is occurring is crucial for future mitigation of the spread of resistance genes as well as their potentially pathogenic hosts (Larsson et al., 2018; Smalla et al., 2018). Our results further stress the need to preferentially use narrow spectrum antibiotics in clinical therapy to maintain a healthy microbiome within the patient that can more easily recover after antibiotic administration (Palleja et al., 2018), thus decreasing the likelihood of positive selection for pathogens that might have acquired resistance when embedded in a community.

## Material and Methods

### Pig faecal community

Pig faeces were collected from four Cornish Black pigs without previous exposure to antibiotics in April 2016 on Healey’s Cornish Cyder farm (Penhallow, Cornwall, United Kingdom). Two hundred grams of faeces from each pig were pooled, mixed with 400ml each of sterile glycerol and 1.8 g/L NaCl solution. The mixture was homogenized for 3 min in a Retsch Knife mill Gm300 (Retsch GmbH, Haan, Germany) at 2000 rotations per minute (rpm), filtered through a sieve (mesh size ~1mm^2^), centrifuged at 500 rpm for 60 s at 4°C and the liquid supernatant fraction was collected and frozen at −80°C as the inoculum.

### Pig fecal extract

Two hundred grams of faeces from each pig were pooled, mixed with 800 mL of sterile 0.9 g/L NaCl solution. The mixture was homogenized for 3 minutes in a Retsch Knife mill Gm300 (Retsch GmbH, Haan, Germany), at 2000 rotations per minute, filtered through a sieve (mesh size ~1 mm^2^) and the liquid fraction was collected. The extract was then centrifuged (3500 rpm, 20 minutes, 4°C), the supernatant collected and autoclaved (121°C, 20 min). The autoclaved extract was centrifuged again (3500 rpm, 20 minutes, 4°C) and the supernatant collected and used as a nutrient supplement.

### Strains

The focal species, *E. coli* MG1655, was chromosomally tagged with a TN7 gene cassette encoding constitutive red fluorescence, expressed by the *mCherry* gene (Remus-Emsermann et al., 2016) to ensure that *E. coli* can be detection and distinguished from other community members after competition based on red fluorescence. The kanamycin resistant, red fluorescent variant containing resistance gene *aph(3′)-IIb* encoding an aminoglycoside 3′-phosphotransferase was created previously (Klümper et al., 2015, 2014).

To create the gentamicin resistant mutant the strain was further tagged through electroporation with the pBAM delivery plasmid containing the mini-TN5 delivery system (Marti-nez-Garci-a et al., 2014; Martínez-García et al., 2011) for gentamicin resistance gene *aacC1* encoding a gentamicin 3′-*N*-acetyltransferase (Kovach et al., 1995). Successful clones were screened for gentamicin resistance (30 μg/mL) and for the chosen clone a single strain growth curve in LB medium was measured to ensure that the cost of the resistance gene was lower than 10% compared to the susceptible strain to ensure competitive ability.

### Competition experiments

Competition experiments as well as initial growth of focal species strains were performed in 25 mL serum flasks with butyl rubber stoppers. As growth medium 10mL of sterile Luria Bertani broth supplemented with 0.1% pig faecal extract, 50 mg/L Cysteine-HCl as an oxygen scrubber and 1 mg/L Resazurin as a redox indicator to ensure anaerobic conditions (Großkopf et al., 2016), was added to each reactor, heated in a water bath to 80°C and bubbled with 100% N_2_ gas until the oxygen indicator Resazurin turned colourless. After cooling down to 37°C the appropriate concentration of antibiotic (AB) was added from a 1000x anaerobic stock solution.

Two isogenic pairs of the focal species, the susceptible, red fluorescent *E. coli* strain with either its gentamicin or kanamycin resistant counterpart, were competed across a gradient of six antibiotic concentrations (Gentamicin [μg/mL]: 0, 0.01, 0.1, 1, 10, 100; Kanamycin [μg/mL]: 0, 0.02, 0.2, 2, 20, 200). Strains as well as the community (100 μL of frozen stock) were grown separately under anaerobic conditions in triplicate reactors, replicates were combined, harvested through centrifugation, washed twice in 0.9% anaerobic NaCl solution and finally resuspended in 0.9% NaCl solution, adjusted to OD_600_ 0.1 (~10^7^ bacteria/mL) and subsequently used in competition experiments. Isogenic strains were mixed at 1:1 ratio (no community treatment), and that mix further added at 10% ratio to 90% of the faecal community (community treatment). Approximately 10^6^ bacteria of either mix were transferred to 6 replicate reactors of each of the antibiotic concentrations and grown at 37°C with 120 rpm shaking for 24h which allowed growth up to carrying capacity. 100μL of each reactor were then transferred to a fresh bioreactor, grown for 24h, transferred for a final growth cycle and finally harvested for subsequent analysis.

### Fitness assay

From each reactor after 3 days (T_3_), as well as the inocula (T_0_), a dilution series in sterile 0.9% NaCl solution was prepared and plated on LB and LB+AB (30 μg/mL Gm or 75 μg/mL Kn). For appropriate dilutions total and resistant red fluorescent *E. coli* colonies were counted under the fluorescence microscope. Plating of the susceptible strain on LB+AB plates further did not lead to any growth of spontaneous mutants. The relative Fitness (*ρ*) of the resistant (r) compared to the susceptible strain (s) strain was subsequently calculated based on their individual growth rate (*γ*) throughout the competition experiment:

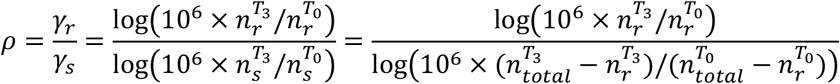

Statistical significant testing (n=6) was performed using a one-tailed t-test against neutral selection (*ρ*=1) and ANOVA corrected for multiple testing to compare the relative fitness of different samples.

### DNA extraction & sequencing

Bacteria from each reactor, as well as inoculum and original pig faecal community were harvested through centrifugation of 2 mL of liquid, followed by DNA extraction using the Qiagen PowerSoil kit as per the manufacturer’s instructions. The quality and quantity of the extractions was confirmed by 1% agarose gel electrophoresis and dsDNA BR (Qubit) respectively.

16S rRNA gene libraries were constructed using multiplex primers designed to amplify the V4 region (Kozich et al., 2013). Amplicons were generated using a high-fidelity polymerase (Kapa 2G Robust), purified with the Agencourt AMPure XP PCR purification system and quantified using a fluorometer (Qubit, Life Technologies, Carlsbad, CA, USA). The purified amplicons were pooled in equimolar concentrations based on Qubit quantification. The resulting amplicon library pool was diluted to 2 nM with sodium hydroxide and 5 mL were transferred into 995mL HT1 (Illumina) to give a final concentration of 10 pM. 600 mL of the diluted library pool was spiked with 10% PhiXControl v3 and placed on ice before loading into Illumina MiSeq cartridge following the manufacturer’s instructions. The sequencing chemistry utilized was MiSeq Reagent Kit v2 (500 cycles) with run metrics of 250 cycles for each paired end read using MiSeq Control Software 2.2.0 and RTA 1.17.28.

Metagenomic libraries were created using the KAPA high throughout Library Prep Kit (Part No: KK8234) optimized for 1ug of input DNA with a size selection and performed with Beckman Coulter XP beads (Part No: A63880). Samples were sheared with a Covaris S2 sonicator (available from Covaris and Life Technologies) to a size of 350bp. The ends of the samples were repaired, the 3′ to 5′ exonuclease activity removed the 3′ overhangs and the polymerase activity filled in the 5′ overhangs creating blunt ends. A single ‘A’ nucleotide was added to the 3’ ends of the blunt fragments to prevent them from ligating to one another during the adapter ligation reaction. A corresponding single ‘T’ nucleotide on the 3’ end of the adapter provided a complementary overhang for ligating the adapter to the fragment ensuring a low rate of chimera formation. Indexing adapters were ligated to the ends of the DNA fragments for hybridisation on a flow cell. The ligated product underwent size selection using the XP beads detailed above, thus removing the majority of un-ligated or hybridized adapters. Prior to hybridisation the samples underwent 6 cycles of PCR to selectively enrich those DNA fragments with adapter molecules on both ends and to amplify the amount of DNA in the library. The PCR was performed with a PCR primer cocktail that anneals to the ends of the adapter. The insert size of the libraries was verified by running an aliquot of the DNA library on a PerkinElmer GX using the High Sensitivity DNA chip (Part No: 5067-4626) and the concentration was determined by using a High Sensitivity Qubit assay. All raw sequencing data has been submitted to ENA under study accession number PRJEB29924.

### 16S Analysis

Sequence analysis was carried out using mothur v.1.32.1 (Schloss et al., 2009) and the MiSeq SOP (Kozich et al., 2013) as accessed on 07.08.2017 on http://www.mothur.org/wiki/MiSeq_SOP. Sequences were classified based on the RDP classifier (Wang et al., 2007). Diversity was assessed based on observed OTUs at 97% sequence similarity. NMDS plots for the community were created after removing all sequences of the focal species *E. coli* based on the Bray-Curtis dissimilarity metric (Bray and Curtis, 1957). Further sample similarity was tested using analysis of molecular variance (AMOVA) a nonparametric analogue of traditional ANOVA testing. AMOVA is commonly used in population genetics to test the hypothesis that genetic diversity between two or more populations is not significantly different from a community created from stochastically pooling these populations (Anderson, 2001; Gravina and Vijg, 2010).

### Metagenomic analysis

Metagenomic samples, as well as a reference genome for the focal species *E. coli* MG1655, were analysed using the ARG-OAP pipeline for antibiotic resistance genes detection from metagenomic data using an integrated structured antibiotic resistance gene database (Yang et al., 2016). This resulted in the abundance of different resistance gene classes and subtypes within these groups normalized by 16S rRNA copy number. Antibiotic resistance genes detected in the *E. coli* reference genome were subtracted from the total number of hits per 16S copy based on the abundance of *E. coli* 16S/total 16S. Further, all antibiotic resistance gene numbers were normalized to the amount of pig faecal community 16S per total 16S copy.

### Antibiotic inhibition testing

To test if any degradation of antibiotics occurred in any of the reactors, 500μL of filter sterilized (0.22 μm^2^ pore size) supernatant of each reactor were applied to a 6mm Grade AA paper disk (Whatman, Maidstone, UK), and a paper disk-agar diffusion assay(Raahave, 1974) was performed on LB medium supplemented with 1% of an overnight culture of the susceptible strain. After 24h incubation at 37°C images of the halo zones were taken using a Leica S8APO stereomicroscope (Leica, Wetzlar, Germany). The area of halo zones was determined by image analysis in Inkscape (version 0.91, http://www.inkscape.org/). Three technical replicate disks for each of the six replicate reactors were averaged, for a total of 18 measurements per concentration.

### Mathematical model

In order to illustrate possible mechanisms underlying the data for bacterial fitness in the presence / absence of the community for varying concentrations of gentamicin and kanamycin, we described our experimental setup mathematically. For this we first developed a discrete-time mathematical model for the growth of the susceptible and drug-resistant bacteria, *s* and *r*, respectively, in the presence or absence of the community, *c*.

#### Bacterial growth

The discrete-time model describing the growth of the bacteria *i*, *i*=*s*,*r*,*c*, is governed by the following iterative model

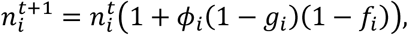

where 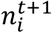 is the size of the population of strain *i* at time *t*+1, and *ϕ*_*i*_ is the maximum growth rate in the absence of competition and drug pressure. The reduction in growth due to density-dependent regulation / resource limitation, given as

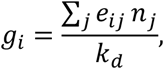

with *k*_*d*_ as the carrying capacity and *e*_*ij*_ being the competition coefficient, describing how much the presence of an allospecific strain *j* impacts the competitive fitness of strain *i*. The reduction in bacterial growth due to drug pressure, *f*_*i*_, is governed by a generalised logistic function

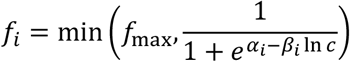

where *c* is the drug concentration (in μg/mL), α_*i*_ and β_*i*_ are the parameters describing the dose-response relationship for strain *i*, and *f*_max_ =0.9 is the maximum growth inhibition.

#### Model simulation and relative fitness calculation

Starting from an initially small number of bacteria in fresh medium, we ran the model for 30 generations, at which point the bacterial population had reached carrying capacity, and diluted the population accordingly. The bacteria were again allowed to grow for 30 generations before being diluted and grown for a final 30 generations. At this point we calculated the relative fitness of the resistant strain as

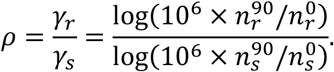

#### Community-dependent change in drug resistance / susceptibility

The kanamycin data seem to suggest that the benefit of the drug resistant bacteria is reduced in the presence of the community at medium to high drug concentrations pointing towards a decrease in the susceptibility of the susceptible strain in a community context. We captured this scenario by making the dose-response parameters *α*_*s*,*r*_ and *β*_*s*,*r*_ explicitly dependent on the density of the community by increasing the resistance of susceptible strain, *s*, i.e.

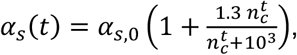

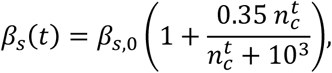

where α_*i*,*0*_ and β_*i*,*0*_ are the time-independent dose-response parameters (Table 2). The effect of density dependence is further illustrated in Figure SI4.

### Parameter estimations

For each drug (gentamicin and kanamycin) we obtained a set of parameter values that resulted in a good overall fit between the model simulations and the data, where the data comprised the observed relative fitness for both sets of experiments (i.e. bacteria grown in the presence and absence of the community) for six different drug concentrations. To allow for logarithmic regression the non-antibiotic control was assumed as one order of magnitude lower than the lowest concentration used in the experiment. The parameter values were determined by minimising the root-mean-square error using an optimisation algorithm akin to simulated annealing (Kirkpatrick et al., 1983). The aim here was not to perform rigorous parameter estimation but rather to find a set of parameters that, given specific model constraints and assumptions, resulted in model behaviours that qualitatively agreed with both the observed dynamics over the repeated growth cycles and the empirically determined fitness values. In fact, our method failed to find a unique set of values that consistently gave the best fitting model, which suggests that the available data was insufficient to determine the global maximum. However, the qualitative relationships between individual parameters and between the parameters comparing the two antimicrobials were fairly consistent between model runs. Tables 1 and 2 list the sets of parameters as used in Figures 1–2.

## Acknowledgments

UK received funding from the European Union’s Horizon 2020 research and innovation program under Marie Skłodowska-Curie grant agreement no. 751699. UK, AB and WG were supported through an MRC/BBSRC grant (MR/N007174/1). XY thanks The University of Hong Kong for a postgraduate studentship.

## Author contributions

UK, LZ, AB and WG conceived the study and designed experiments; UK performed experimental work; MR performed mathematical modelling; UK, XY and TZ performed sequencing analysis; UK, AB, WG analysed data and wrote the manuscript.

## Competing interests

The authors declare no competing interests.

## Materials & Correspondence

All correspondence and material requests should be addressed to UK.

