## Supplemental Figure 1-4 for "Selection for antibiotic resistance is reduced when embedded in a natural microbial community"

### Title

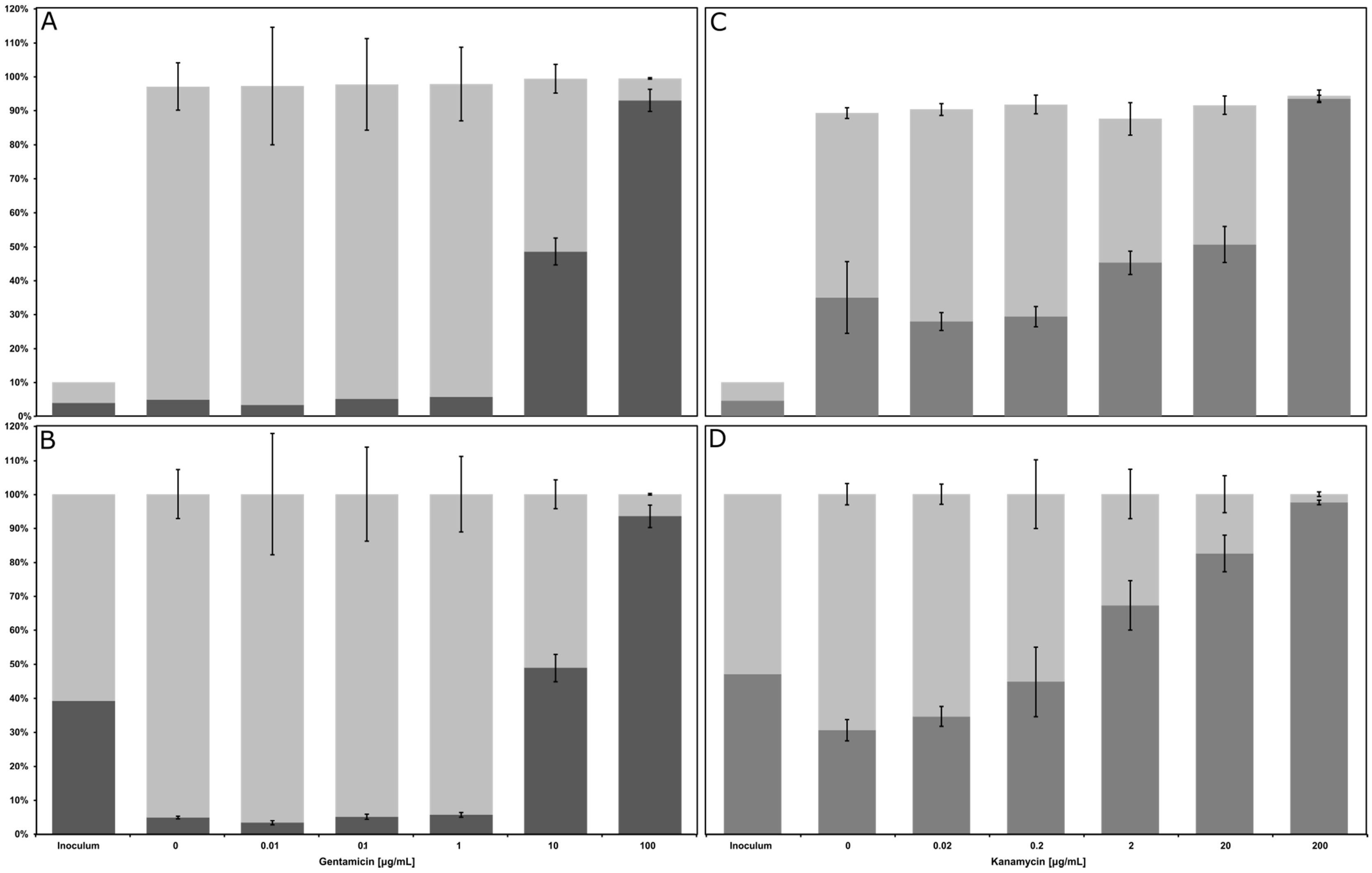


**Figure SI1: Relative abundance of the focal species after 3 days of the competition experiment**

Values shown are mean ± SD (n=6, resistant: black, susceptible: grey). (A) gentamicin, absolute values including community; (B) gentamicin, ratio of the isogenic pair of the focal species (C) kanamycin, absolute values including community; (D) kanamycin, ratio of the isogenic pair of the focal species.

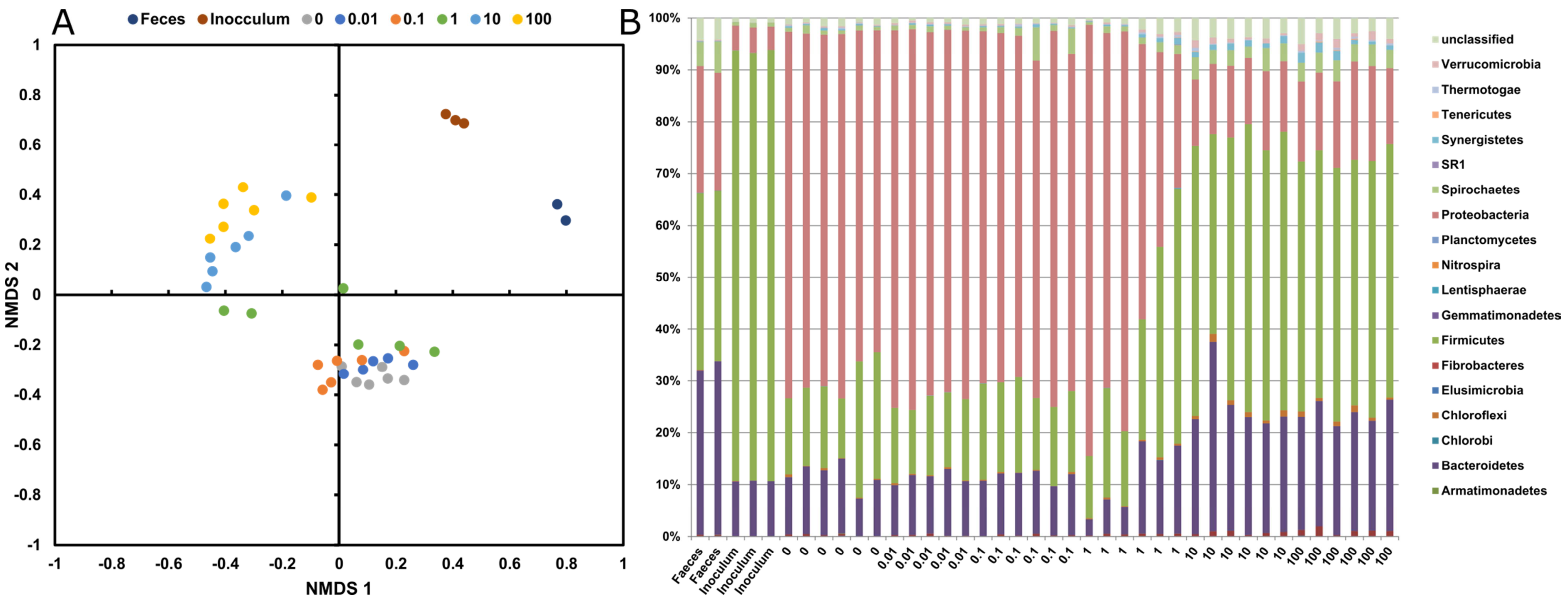


**Figure SI2. Community analysis for gentamicin reactors.**

(A): Non-metric 2-dimensional scaling analysis (NMDS) revealing distinct clustering of original fecal community, inoculum and reactors after 3 day incubation. Ordination based on the Bray-Curtis dissimilarity metric. (B) Bar chart based on phylum distribution.

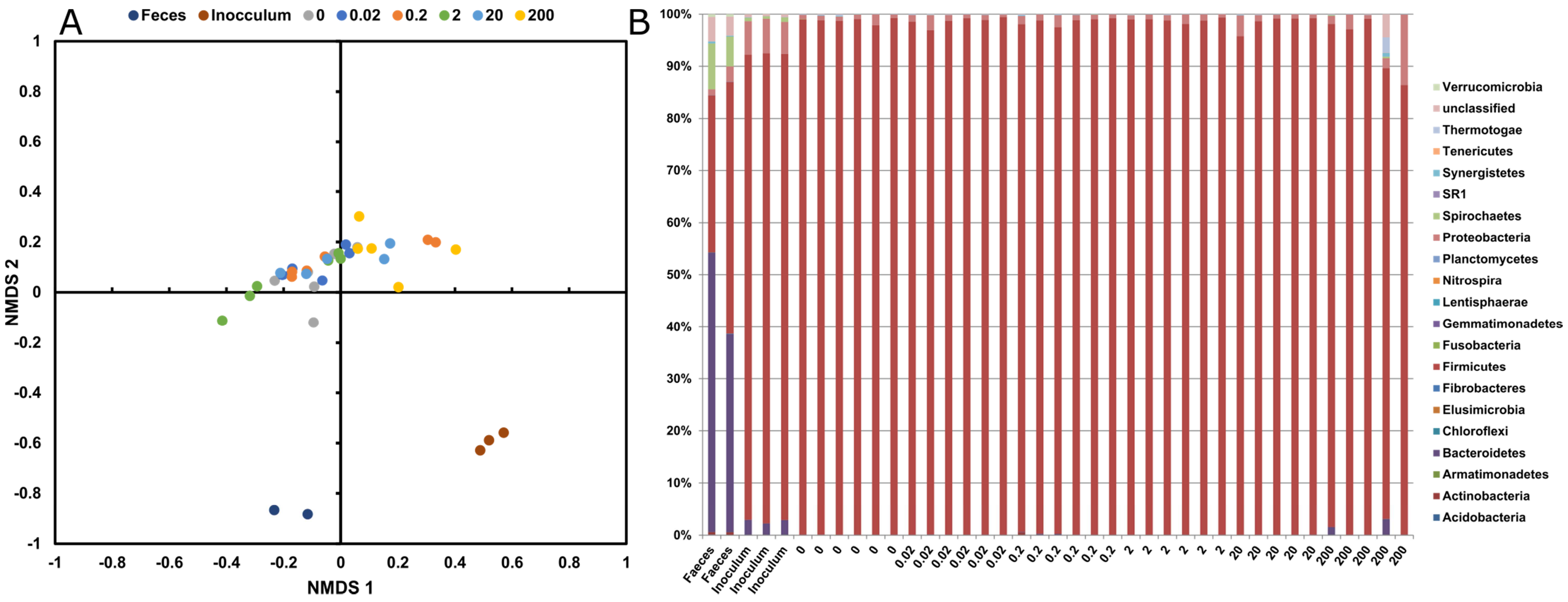


**Figure SI3. Community analysis for kanamycin reactors.**

(A) Non-metric 2-dimensional scaling analysis (NMDS) revealing distinct clustering of original fecal community, inoculum and reactors after 3 day incubation. Ordination based on the Bray-Curtis dissimilarity metric. (B) Bar chart based on phylum distribution.

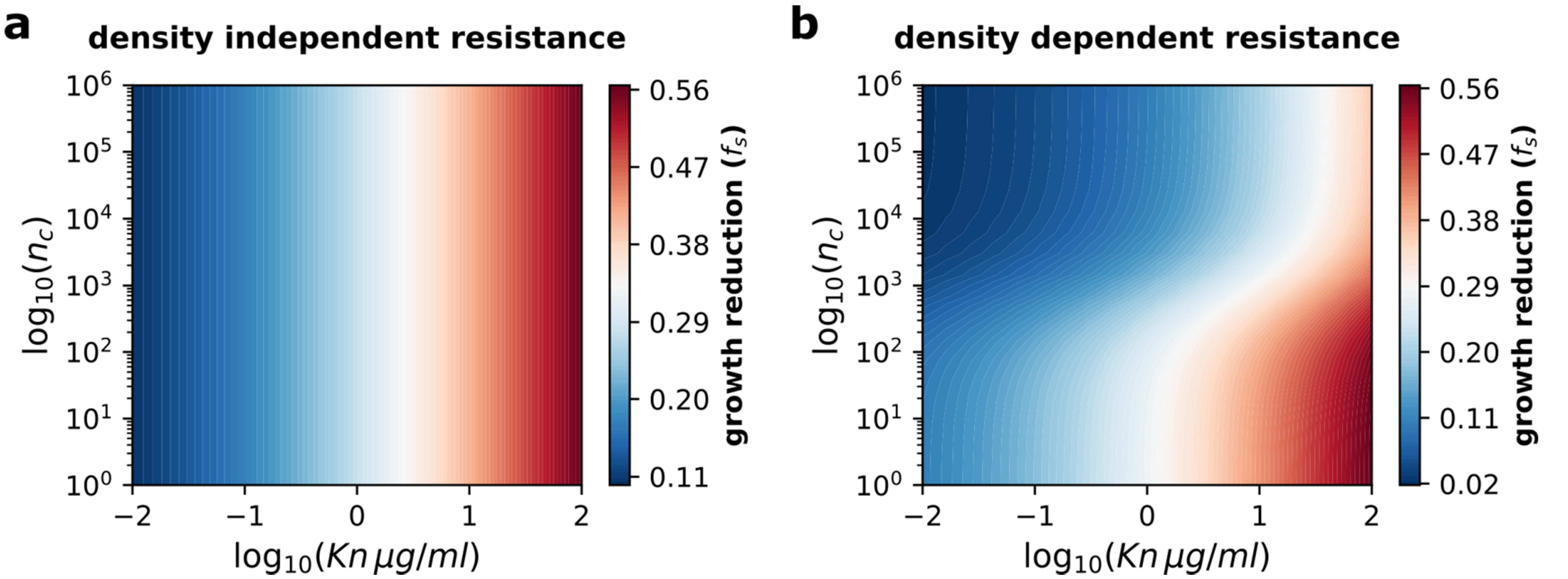


**Figure SI4. Contour plots showing the growth reduction of the susceptible strain**,

dependent on the Kanamycin concentration and size of the community, for the two scenarios used in the mathematical model: density independent (a) and dependent (b) resistance
